# Sequential Image Classification of Human-Robot Walking Environments using Temporal Neural Networks

**DOI:** 10.1101/2023.11.10.566555

**Authors:** Bogdan Ivanyuk-Skulskiy, Andrew Garrett Kurbis, Alex Mihailidis, Brokoslaw Laschowski

## Abstract

Robotic prosthetic legs and exoskeletons require real-time and accurate estimation of the walking environment for smooth transitions between different locomotion mode controllers. However, previous studies have mainly been limited to static image classification, therein ignoring the temporal dynamics of human-robot locomotion. Motivated by these limitations, here we developed several state-of-the-art temporal convolutional neural networks (CNNs) to compare the performances between static vs. sequential image classification of real-world walking environments (i.e., level-ground terrain, incline stairs, and transitions to and from stairs). Using our large-scale image dataset, we trained a number of encoder networks such as VGG, MobileNetV2, ViT, and MobileViT, each coupled with a temporal long short-term memory (LSTM) backbone. We also trained MoViNet, a new video classification model designed for mobile and embedded devices, to further compare the performances between 2D and 3D temporal deep learning models. Our 3D network outperformed all the hybrid 2D encoders with LSTM backbones and the 2D CNN baseline model in terms of classification accuracy, suggesting that network architecture can play an important role in performance. However, although our 3D neural network achieved the highest classification accuracy, it had disproportionally higher computational and memory storage requirements, which can be disadvantageous for real-time control of robotic leg prostheses and exoskeletons with limited onboard resources.

## I. Introduction

Robotic leg prostheses and exoskeletons require accurate and real-time state estimation to transition between different controllers, especially over complex terrains such as stairs [1]. Cameras have been used for environment state estimation in combination with inertial sensors, used for locomotor state estimation [2]. Early systems for environment recognition were limited to statistical pattern recognition and machine learning algorithms that required manual feature engineering and/or were developed using relatively small image datasets [3]-[11]. Recent studies [12]-[19] have focused on using deep learning and large-scale datasets, such as ExoNet [20] and StairNet [21], to develop systems that can generalize to diverse walking environments. These systems use convolutional neural networks (CNNs) and transfer learning for image classification such that the model weights are trained on large datasets like ImageNet [22] and fine-tuned on downstream tasks with modified head layers. However, these systems classify each frame independently, therein ignoring the temporal dynamics of human-robot locomotion, which motivated our study.

Research in autonomous driving has shown that sequential data rather than independent frames, comparable to biological vision systems, can improve performance for steering wheel angle prediction during self-driving [23]. Taking inspiration from humans and autonomous cars, here we developed and trained several state-of-the-art temporal neural networks on a large-scale dataset to compare the performances between using static vs. sequential data for image classification of real-world walking environments, with an emphasis on stair recognition. We focus on stairs because it presents a large safety critical application for robotic leg control.

In this study, we quantitatively compared the performances between static vs. sequential image classification through the development and evaluation of several state-of-the-art temporal neural networks, specifically a 3D CNN model designed for temporal feature learning and hybrid models with 2D encoders paired with temporal backbones. We analyzed the performance of our temporal models and compared the results to a static CNN baseline model [12] in terms of classification accuracy, inference speed, and efficiency [24].

## II. Methods

### A. Image Dataset

To train and test our image classification models, we used the open-source StairNet dataset [21] with four environment classes: level-ground terrain (LG), level-ground transition to incline stairs (LG-IS), incline stairs (IS), and incline stairs transition to level-ground (IS-LG). The StairNet dataset includes videos recorded in urban environments using a wearable camera. Video frames were recorded with a step size of 6, with each video having a variable number of episodes (i.e., continuous frames of stair climbing environments) ranging from 43 to 687, where each episode has 20 frames on average (Table 1). The StairNet dataset contains unbalanced classes due to the different frequencies of real-world walking environments. The dataset mainly comprises level-ground (86% of samples) and incline stairs (9%). It has two minority classes, IS-LG and LGIS, which contain roughly 2% and 3% of the total samples, respectively. This imbalance is important when selecting our classification and resampling techniques.

**Table 1.** The class distributions for the training, validation, and test dataset splits used to develop our deep learning models.

| Split | Number of videos | Number of frames | Class | Percent of class across splits (%) |
| --- | --- | --- | --- | --- |
| Train | 45 | 466,177 | IS | 84.5 |
|  |  |  | ISLG | 83.4 |
|  |  |  | LG | 82.5 |
|  |  |  | LGIS | 80.0 |
| Validation | 6 | 32,487 | IS | 5.3 |
|  |  |  | ISLG | 6.5 |
|  |  |  | LG | 6.3 |
|  |  |  | LGIS | 7.5 |
| Test | 6 | 56,729 | IS | 10.2 |
|  |  |  | ISLG | 10.1 |
|  |  |  | LG | 11.2 |
|  |  |  | LGIS | 12.5 |

Previous studies using StairNet dataset for deep learning [12]-[14] have used a train-validation-test split that randomly selects single frames throughout the dataset. This validation method can be fragile to data leakage for the test set as it contains similar neighbouring frames in both the model training and evaluation. To prevent data leakage, we used a video-based validation method. This method includes assigning all frames within a video episode (i.e., a group of neighbouring frames) to one of the train-validation-test splits, thus preventing overlap in the splits and resulting in a better estimation of the real-world performance and generalizability [25]. The class distribution in each split was preserved to match that of previous research [12] (Table 1).

### B. Image Processing

We divided each video into clips comprising several frame sequences. Each video frame sequence includes a fixed anchor frame with a corresponding class label and an *n* number of neighbouring frames sequenced in temporal order. We selected the *n* for each video frame sequence length to be 5. Since each frame in the StairNet dataset represents every sixth frame in the original ExoNet video sequences, extracting five frames per sample provides a historical context of ∼1 second with a recording speed of 24 frames per second. Given that each frame sequence within the video clip has its own label, two training settings were used. The first training setting was sequence-to-sequence classification in which the model predicted the labels for each frame of the video clip. The second training setting was a sequence-to-one classification, where the sequence label corresponded to the anchor frame label.

Most of the data processing was performed before model training because of the large dataset size and corresponding data-loading speed constraints. Data preprocessing included normalizing all frames to be in [0, 1] range, resizing to match the deep learning model input size of 256 × 256 in width and height, with an additional crop to 224 × 224. Once the data preprocessing was complete, we created a TFRecords file with all training and test data to allow for efficient data loading for the TensorFlow framework using the TFRecords series of binary records. We also achieved similar data loading and training times for the PyTorch framework using the lightning memory-mapped database.

### C. Temporal Neural Networks

We experimented with several temporal neural networks for sequential image classification (Fig. 1). We started by experimenting with different model architectures, including MoViNet [26], a lightweight 3D CNN architecture, and several encoder architectures, including VGG-19 [27], MobileNetV2 [28], MobileViT [29], and ViT-B16 [30], each paired with a temporal long-short term memory (LSTM) backbone [31] or transformer encoder [32].

**Fig. 1.**
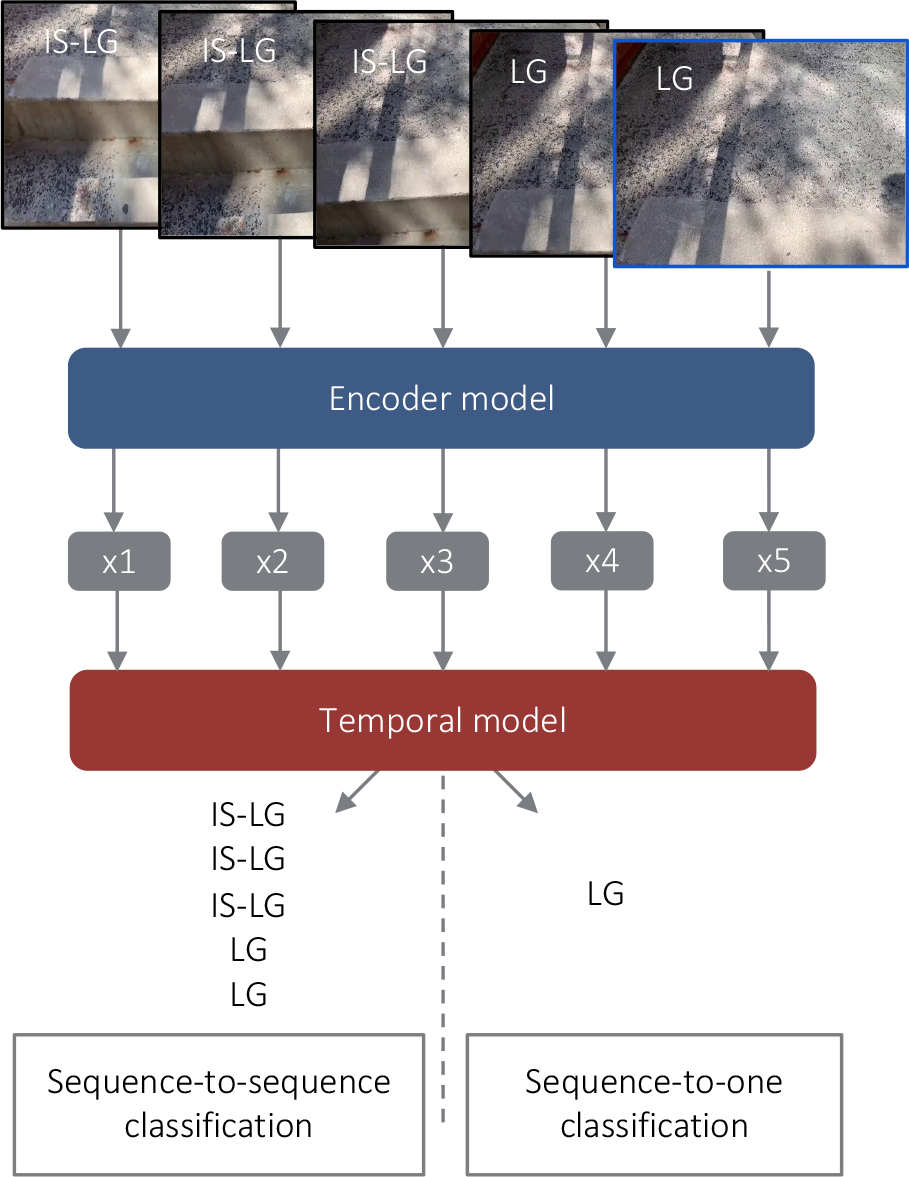
Schematic of the temporal neural networks used for sequential image classification of human-robot walking environments with the two variants of prediction methods: sequence-to-sequence and sequence-to-one classification.

Our model selection process comprised two stages. In the first stage, we assessed model statistics, taking into account factors such as the architecture size and the number of FLOPs. Large models, such as VGG, ViT, and models with temporal transformer encoder blocks, were excluded during this stage. In the second stage, we conducted a brief training session of 5 epochs for each model. This preliminary training allowed us to identify the top five best-performing models: 3D MoViNet, and MobileViT and MobileNetV2 as encoders, coupled with LSTM or transformer blocks as temporal classifiers. We then performed an extensive hyperparameter search and conducted more extensive training.

#### 3D-CNN

The first model we experimented with was MoViNet, a modified MobileNetV3 [33] architecture adapted to videos. We used a stream buffer to minimize model memory growth, which is a cache feature that is applied to the boundaries of the video sub-sequences and mitigates the growth of the model memory in proportion to the number of frames in the sequence. Initially, the cache is zero-initialized. The feature map is then computed by applying a temporal operation (i.e., 3D convolution) over the concatenation of the buffer and subsequence. In later feature maps, the buffer is updated by concatenating the current buffer and new sequence using the following formula:

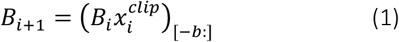

where *B*_*i*_ is the buffer, 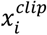 is the original input sequence, and [−*b*] is the selection of the last *b* frames of the concatenated feature sequence. The stream buffer reduces the memory usage of the MoViNet model at the cost of a small accuracy degradation, which we mitigated using an ensemble of models. We trained two identical MoViNet architectures with a half-frame rate to achieve this. During inference, the input sequence was fitted to both networks, and the mean value of the two models was obtained and passed through the softmax classification activation function.

#### 2D-CNN with Temporal Blocks

We then experimented with a hybrid MobileNetV2 architecture, selected for its efficient model design, optimized for mobile and embedded computing devices, coupled with an LSTM temporal layer. The encoder architecture comprises an input convolutional layer and 19 residual bottleneck layers. Each residual bottleneck layer consists of three parts: an expansion layer that projects the input tensor to a higher-dimensional space where the number of channels is increased according to a hyperparameter *t* called the expansion factor [28], a depth-wise convolution with ReLU that decreases the size of the feature map but preserves the number of channels, and a 1 × 1 convolution layer that projects the feature tensor to a lower-dimension tensor by decreasing the number of channels. MobileNetV2 was applied to each sequence frame separately, resulting in a stack of feature maps. This stack was then passed through the LSTM layer to capture temporal dynamics. The output of the LSTM layer was either a sequence of labels for sequence-to-sequence classification or the last predicted label on the recurrency operation of the LSTM layer for sequence-to-one classification. Although the latter approach may appear to be more intuitive, the former suggests that sequence-to-sequence classification can yield a more intricate understanding of temporal variations in terrain.

#### Hybrid ViT with Temporal Blocks

Finally, we experimented with MobileViT, a hybrid encoder model that combines local information from convolutional layers and global information from feature maps extracted using MobileViT blocks in the later layers. The convolution layers are represented by a strided 3 × 3 convolution on the input and several MobileNetV2 blocks. MobileViT applies standard convolution to encode local spatial information and point-wise convolution to create a projection to a high-dimensional space. These high-dimensional projections are unfolded into *N* non-overlapping flattened patches and encoded using transformer blocks. The transformer outputs are then projected back to the original low-dimensional space and fused with the original feature maps. Similar to MobileNetV2, the MobileViT model was applied to each sequence frame separately, resulting in a sequence of feature maps corresponding to the number of frames in the sample. These feature maps were passed through the LSTM layer to capture the temporal dynamics of the sequence feature maps. In sequence-to-sequence classification, the output of the last transformer block is passed through a linear classification head. In sequence-to-one classification, we flatten the transformer layer’s output before the classification head.

### D. Hyperparameter Optimization

We conducted hyperparameter optimization using the KerasTuner to fine-tune the configuration of our top-5 best-performing models, derived from our initial experimentation. These models collectively shared a set of common hyperparameters, which include dropout rate and the number of trainable layers. Furthermore, each model contained model-specific hyperparameters tailored to its architecture. For example, the MoViNet search space optimized the activation function, convolutional filter width, and the number of feature map channels. The MobileNetV2 with LSTM optimized the MobileNet block expansion rate, bias term, the quantity of LSTM units in each layer, and the number of LSTM layers. Similarly, the MobileViT with transformer optimized the MobileNet block expansion rate, the number of attention heads within the transformer, the number of dense units within the transformer block, and the dimensionality of the dense layers. For general optimization, all models were also tuned for learning rate, learning rate scheduler, and choice of optimizer.

After finding the optimal hyperparameters, we trained each model for 20 epochs using an NVIDIA Tesla V100 32GB GPU. We used a learning rate of 0.0001 and the Adam [34] optimizer with a cosine annealing learning rate scheduler, as these hyperparameters presented the best results on the initial hyperparameter selection stage.

### E. Performance Evaluation

To quantitatively evaluate and compare the performance of each temporal neural network, we used classification accuracy, precision, recall, F1-score, and model characteristics, including the number of floating-point operations (FLOPS) and the number of frames processed per second. However, our main metric for comparison was NetScore [24], which balances the model performance with efficiency and is represented by the following equation:

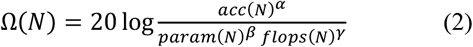

where *acc*(*N*) is the classification accuracy (%), *param*(*N*) is the number of model parameters, which is indicative of the memory storage requirements, *flops*(*N*) is the number of floating point operations, which is indicative of the computing requirements, and *α, β, γ* are coefficients that control the influence of each parameter on the overall NetScore.

Sequence-to-one models were evaluated as regular classification models, where the predicted label of the sequence is compared to the ground truth. Sequence-to-sequence models were evaluated in two settings. The first setting was sequence-to-sequence evaluation, where the predicted sequence of labels was compared to the ground truth sequence of labels. The second setting was sequence-to-one evaluation, where only the anchor frame label is selected for comparison. The later approach is used to compare the performance of sequence-to-sequence models to sequence-to-one models.

## III. Results

Our experimental results are summarized in Fig. 2 and Table 2. The 3D-CNN MoViNet model achieved the highest classification accuracy and F1-score of 98.3% and 98.2%, respectively. The other hybrid models, including 2D-CNN encoder and temporal blocks (i.e., MobileNetV2 with LSTM and MobileViT with LSTM), struggled to capture inter-frame dependencies using minimal sequences (i.e., five frames per sample) [35] and achieved a lower classification performance than our 3D network. Of the temporal models that we studied, the 3D MoViNet network had the best NetScore of 167.4, out-performing the hybrid 2D encoder models with scores of 154.9 and 132.1 for MobileViT with LSTM and MobileNetV2 with LSTM, respectively. However, due to the relatively low number of parameters and numerical operations, our 2D-CNN baseline model achieved the overall best NetScore of 186.8.

**Table 2.** The experimental results for our six different deep learning models, including the baseline 2D MobileNetV2 model, the temporal sequence-to-one models (i.e., MoViNet, MobileViT-XXS + LSTM, and MobileNetV2 + LSTM), and the temporal sequence-to-sequence models (i.e., MobileNetV2 + LSTM with sequence-to-sequence evaluation and MobileNetV2 + LSTM with sequence-to-one evaluation). The best values for each metric are shown in bold.

| Model | Classification Type | Parameters (millions) | FLOPs (billions) | Accuracy (%) | F1-score (%) | NetScore | Frames per second |
| --- | --- | --- | --- | --- | --- | --- | --- |
| MoViNet | Seq-to-one | 4.03 | 2.5 | **98.3** | **98.2** | 167.4 | 13.5 |
| MobileViT-LSTM | Seq-to-one | 3.36 | 9.84 | 97.0 | 96.8 | 154.9 | 17.6 |
| MobileNetV2-LSTM | Seq-to-one | 6.08 | 53.96 | 97.3 | 97.0 | 132.1 | 17.7 |
| MobileNetV2-LSTM | Seq-to-seq, seq-to-seq | 5.93 | 50.97 | 70.7 | 79.9 | 120.2 | 17.6 |
| MobileNetV2-LSTM | Seq-to-seq, seq-to-one | 5.93 | 50.97 | 43.2 | 53.8 | 100.5 | 17.6 |
| Baseline model | Single input/output | **2.26** | **0.61** | 97.2 | 97.2 | **186.8** | **22.6** |

**Fig. 2.**
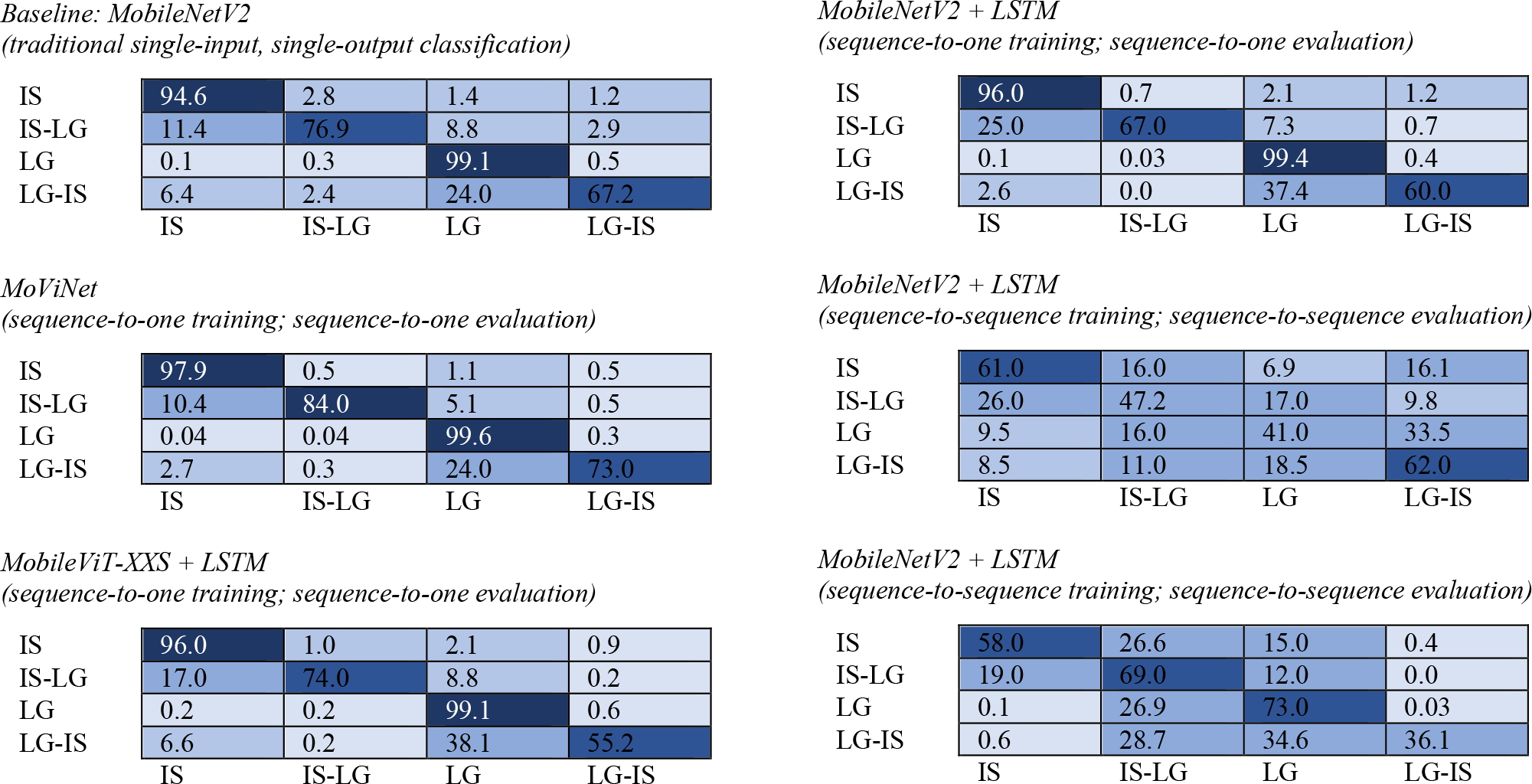
The normalized confusion matrices during inference for our six deep learning models, including the baseline 2D MobileNetV2 model, the temporal sequence-to-one models (i.e., MoViNet, MobileViT-XXS + LSTM, and MobileNetV2 + LSTM), and the temporal sequence-to-sequence models (i.e., MobileNetV2 + LSTM with sequence-to-sequence evaluation and MobileNetV2 + LSTM with sequence-to-one evaluation).

Our MobileViT with LSTM model (i.e., the hybrid of a CNN and transformer blocks) had slightly lower classification performance compared to MobileNetV2 with LSTM with F1-scores of 96.8% and 97.0%, respectively. However, MobileViT has disproportionally less model parameters (3.4 million) and numerical operations (9.8 billion FLOPS) compared to 6.1 million parameters and 54 billion FLOPS for MobileNetV2, resulting in a better NetScore. We also studied the impact of resampling input frames to produce a prediction sequence (i.e., sequence-to-sequence) instead of outputting a single prediction for the sequence without resampling the frames beyond the anchor (i.e., sequence-to-one). Our findings showed an increase in performance using sequence-to-one classification, with an accuracy of 97.3% compared with 70.7% using sequence-to-sequence.

## IV. Discussion

In this study, we developed a number of state-of-the-art temporal neural networks to quantitatively compare the performances between static vs. sequential image classification of human-robot walking environments, with an emphasis on stair recognition. We used the open-source StairNet dataset to train and test our deep learning models. We experimented with different architectures, including a 3D-CNN model and several hybrid 2D encoder models (i.e., VGG, MobileNetV2, ViT, and MobileViT) each paired with a temporal backbone of LSTM or transformer encoder blocks. Of the temporal networks that we studied, the 3D video classification model achieved the highest accuracy of 98.3%, outperforming our 2D-CNN baseline model with an accuracy of 97.2%. However, our 3D model had significantly more parameters (4 million) compared to 2.3 million of the 2D-CNN model, presenting a tradeoff between size and accuracy, which is important for onboard real-time inference with embedded systems used in robotic leg prostheses and exoskeletons.

Unlike other visual perception systems for human-robot walking [3]-[11], our models offer several advantages. We developed deep learning models that use a multilayer neural network on sequences of frames to automatically learn and extract optimal video features from training data, eliminating the need of classical machine learning that requires time-consuming hand engineering. Our models are also trained using StairNet, which is currently the largest open-source image dataset of real-world stair environments. The dataset consists of over 515,000 labelled images, a substantial increase in size compared to previous research such as the convolutional neural network by [4] that was developed using only 34,000 labelled images.

Unlike many previous studies [12]-[18], we uniquely studied sequential image classification, which takes advantage of the cyclical nature of walking and uses temporal relations in the observed environment. This contribution allowed us to exceed the previous state-of-the-art [12], whereby our 3D-CNN model (98.3% accuracy) outperformed the original StairNet model (97.2% accuracy) when evaluated on the same dataset using our video-based validation method. These findings suggest that using temporal data rather than independent images can improve the visual perception accuracy of human-robot walking environments, aligning with research findings in autonomous driving [23].

Our method for video splits for the StairNet dataset also provides a more accurate ‘true’ estimate for generalization to real-world stair recognition. Previous validation methods used a train-validation-test split that randomly select single frames throughout the dataset. This can cause data leakage for the test set since similar neighbouring frames may be included in both the model training and evaluation. Our method for data splits assigns all frames within a video episode (i.e., a group of neighbouring frames) to a single training, validation, or test set to prevent overlap and data leakage between splits.

Despite this progress, our study has several limitations. All our classification models require processing each frame for the encoder and temporal blocks separately. This results in a computational cost of *t* ∗ *O*(*encoder model complexity*), where *t* is the number of frames in each sample. This computational cost limits efficiency and usability when deployed on mobile and embedded devices for real-world applications. Recent studies in transformers [36] and multilayer perceptrons [37] address this limitation by adapting the models to take 3D sequence inputs by modifying the patch-embedding block. The tunnels of frame sequences are treated as separate tokens and linearly projected to match the input requirements of the model. This method in combination with efficient transformer architectures, such as reversible vision transformers [38] or EdgeViT [39], seems promising and a potential direction for future research.

Additionally, we were only able to make limited comparisons between the performances of our system and previous research since we used a novel method for the data splits for the StairNet dataset, which, combined with temporal classification, significantly impacts accuracy. To compare our temporal results with the current state-of-the-art that used static images [12], we re-evaluated the model using our novel validation method. We aim to promote direct comparisons in the future by making our StairNet dataset manipulation and preprocessing methods publicly available to the research community.

In summary, here we showed that, compared to static image classification using 2D convolutional neural networks, using temporal neural networks with sequential inputs can improve the classification accuracy of real-world walking environments, with an emphasis on stair recognition. However, these improvements in performance come at the cost of greater computational and memory storage requirements, which can be disadvantageous for robotic leg prostheses and exoskeletons with limited onboard resources. For future research, we plan to test and optimize the inference speed of our deep learning models when deployed on embedded devices, such as those used in smart glasses [40], and data fusion with other sensors like surface electromyography for robotic leg control.

## Acknowledgment

We want to thank members of the Bionics Lab, part of the Artificial Intelligence and Robotics in Rehabilitation Team at the KITE Research Institute, Toronto Rehabilitation Institute, for their support. This research is dedicated to the people of Ukraine in response to the 2022 Russian invasion.

## References

[1] M. Grimmer, R. Riener, C. J. Walsh, and A. Seyfarth, “Mobility related physical and functional losses due to aging and disease - A motivation for lower limb exoskeletons,” Journal of NeuroEngineering and Rehabilitation, Jan. 2019.

[2] O. Tsepa, R. Burakov, B. Laschowski and A. Mihailidis, “Continuous prediction of leg kinematics during walking using inertial sensors, smart glasses, and embedded computing,” IEEE International Conference on Robotics and Automation (ICRA), May 2023.

[3] N. E. Krausz and L. J. Hargrove, “Recognition of ascending stairs from 2D images for control of powered lower limb prostheses,” IEEE/EMBS Conference on Neural Engineering (NER), Apr. 2015.

[4] B. Laschowski, W. McNally, A. Wong, and J. McPhee, “Preliminary design of an environment recognition system for controlling robotic lower-limb prostheses and exoskeletons,” IEEE International Conference on Rehabilitation Robotics (ICORR), Jun. 2019.

[5] G. Khademi and D. Simon, “Convolutional neural networks for environmentally aware locomotion mode recognition of lower-limb amputees,” ASME Dynamic Systems and Control Conference (DSCC), Nov. 2019.

[6] N. E. Krausz, T. Lenzi, and L. J. Hargrove, “Depth sensing for improved control of lower limb prostheses,” IEEE Transactions Biomedical Engineering, Nov. 2015.

[7] Y. Massalin, M. Abdrakhmanova, and H. A. Varol, “User-independent intent recognition for lower limb prostheses using depth sensing,” IEEE Transactions Biomedical Engineering, Aug. 2018.

[8] H. A. Varol and Y. Massalin, “A feasibility study of depth image based intent recognition for lower limb prostheses,” IEEE Engineering in Medicine and Biology Society (EMBC), Aug. 2016.

[9] B. Zhong, R. L. da Silva, M. Li, H. Huang, and E. Lobaton, “Environmental context prediction for lower limb prostheses with uncertainty quantification,” IEEE Transactions on Automation Science and Engineering, Apr. 2021.

[10] B. Zhong, R. L. da Silva, M. Tran, H. Huang, and E. Lobaton, “Efficient environmental context prediction for lower limb prostheses,” IEEE Transactions on Systems, Man, and Cybernetics, Jun. 2022.

[11] K. Zhang et al., “A subvision system for enhancing the environmental adaptability of the powered transfemoral prosthesis,” IEEE Transactions on Cybernetics, Jun. 2021.

[12] A. G. Kurbis, B. Laschowski, and A. Mihailidis, “Stair recognition for robotic exoskeleton control using computer vision and deep learning,” International Conference on Rehabilitation Robotics (ICORR), Jul. 2022.

[13] A. G. Kurbis, A. Mihailidis, and B. Laschowski, “Development and mobile deployment of a stair recognition system for human-robot locomotion.” bioRxiv, Apr. 2023.

[14] D. Kuzmenko, O. Tsepa, A. G. Kurbis, A. Mihailidis, and B. Laschowski, “Efficient visual perception of human-robot walking environments using semi-supervised learning.” bioRxiv, Jun. 2023.

[15] B. Laschowski, W. McNally, A. Wong, and J. McPhee, “Environment classification for robotic leg prostheses and exoskeletons using deep convolutional neural networks,” Frontiers in Neurorobotics, Feb. 2022.

[16] J. Xue, H. Zhang and K. Dana, “Deep texture manifold for ground terrain recognition,” IEEE/CVF Conference on Computer Vision and Pattern Recognition (CVPR), Jun. 2018

[17] V. Suryamurthy, et al. “Terrain segmentation and roughness estimation using RGB data: Path planning application on the CENTAURO robot,” IEEE International Conference on Humanoid Robots, Oct. 2019

[18] E. Tricomi et al., “Environment-based assistance modulation for a hip exosuit via computer vision”, IEEE Robotics and Automation Letters, May 2023

[19] A. H. A. Al-Dabbagh and R. Ronsse, “Depth vision-based terrain detection algorithm during human locomotion,” IEEE Transactions on Medical Robotics and Bionics, Nov. 2022

[20] B. Laschowski, W. McNally, A. Wong, and J. McPhee, “ExoNet database: Wearable camera images of human locomotion environments,” Frontiers in Robotics and AI, Dec. 2020.

[21] A.G. Kurbis, D. Kuzmenko, B. Ivanyuk-Skulskiy, A. Mihailidis, B. Laschowski, “StairNet: Visual recognition of stairs for human-robot locomotion,” arXiv, Oct. 2023.

[22] J. Deng, W. Dong, R. Socher, L.-J. Li, K. Li, and L. Fei-Fei, “ImageNet: A large-scale hierarchical image database,” IEEE/CVF Conference on Computer Vision and Pattern Recognition (CVPR), Jun. 2009.

[23] H. M. Eraqi, M. N. Moustafa, and J. Honer, “End-to-end deep learning for steering autonomous vehicles considering temporal dependencies,” arXiv, Nov. 2017.

[24] A. Wong, “NetScore: Towards universal metrics for large-scale performance analysis of deep neural networks for practical on-device edge usage,” arXiv, Aug. 2018.

[25] Y. LeCun, Y. Bengio, and G. Hinton, “Deep learning,” Nature, May 2015.

[26] D. Kondratyuk et al., “MoViNets: Mobile video networks for efficient video recognition,” IEEE/CVF Conference on Computer Vision and Pattern Recognition (CVPR), Jun. 2021.

[27] K. Simonyan and A. Zisserman, “Very deep convolutional networks for large-scale image recognition,” arXiv, Apr. 2015.

[28] A. G. Howard et al., “MobileNets: Efficient convolutional neural networks for mobile vision applications.” arXiv, Apr. 2017.

[29] Mehta, Sachin, and Mohammad Rastegari, “MobileVit: light-weight, general-purpose, and mobile-friendly vision transformer,” arXiv, Oct. 2021.

[30] A. Dosovitskiy et al., “An image is worth 16×16 words: Transformers for image recognition at scale,” arXiv, Jun. 2021.

[31] S. Hochreiter and J. Schmidhuber, “Long short-term memory,” Neural Computation, Nov. 1997.

[32] A. Vaswani et al., “Attention is all you need,” arXiv, Jun. 2017.

[33] A. Howard et al., “Searching for MobileNetV3,” IEEE/CVF International Conference on Computer Vision (ICCV), Oct. 2019.

[34] D. P. Kingma and J. Ba, “Adam: A method for stochastic optimization,” arXiv, Jan. 2017.

[35] J. Carreira and A. Zisserman, “Quo vadis, action recognition? A new model and the kinetics dataset,” IEEE/CVF Conference on Computer Vision and Pattern Recognition (CVPR), Jul. 2017.

[36] Z. Liu et al., “Video swin transformer,” IEEE/CVF Conference on Computer Vision and Pattern Recognition (CVPR), Jun. 2022.

[37] D. J. Zhang et al., “MorphMLP: An efficient MLP-like backbone for spatial-temporal representation learning,” European Conference on Computer Vision (ECCV), Oct. 2022.

[38] K. Mangalam et al., “Reversible vision transformers,” IEEE/CVF Conference on Computer Vision and Pattern Recognition (CVPR), Jun. 2022.

[39] J. Pan et al., “EdgeViTs: Competing light-weight CNNs on mobile devices with vision transformers,” European Conference on Computer Vision (ECCV), Oct. 2022.

[40] D. Rossos, A. Mihailidis, B. Laschowski, “AI-powered smart glasses for sensing and recognition of human-robot walking environments,” bioRxiv, Oct. 2023.

